# High Expression of Glycolytic Genes in Clinical Glioblastoma Patients Correlates with Lower Survival

**DOI:** 10.1101/2021.10.07.463555

**Authors:** Kimberly M Stanke, Carrick Wilson, Srivatsan Kidambi

## Abstract

Glioblastoma (GBM), the most aggressive brain tumor, is associated with a median survival at diagnosis of 16-20 months and limited treatment options. The key hallmark of GBM is altered tumor metabolism and marked increase in the rate of glycolysis. Aerobic glycolysis along with elevated glucose consumption and lactate production supports rapid cell proliferation and GBM growth. In this study, we examined the gene expression profile of metabolic targets in GBM samples from patients with low grade glioma (LGG) and GBM. We found that gene expression of glycolytic enzymes is up-regulated in GBM samples and significantly associated with an elevated risk for developing GBM. Our findings of clinical outcomes showed that GBM patients with high expression of HK2 and PKM2 in the glycolysis related genes and low expression of genes involved in mitochondrial metabolism-SDHB and COX5A related to tricarboxylic acid (TCA) cycle and oxidative phosphorylation (OXPHOS), respectively, was associated with poor patient overall survival. Surprisingly, expression levels of genes involved in mitochondrial oxidative metabolism are markedly increased in GBM compared to LGG but was lower compared to normal brain. The fact that in GBM the expression levels of TCA cycle and OXPHOS-related genes are higher than those in LGG patients suggests the metabolic shift in GBM cells when progressing from LGG to GBM. These results are an important step forward in our understanding of the role of metabolic reprogramming in glioma as drivers of the tumor and could be potential prognostic targets in GBM therapies.

## 1. INTRODUCTION

Glioblastoma Multiforme (GBM) is the most prolific and deadly malignant brain tumor in the United States with an age-adjusted incidence rate of 3.2 cases per 100,000 population and more than 13,000 cases projected in 2020 [1; 2; 3]. The median survival at GBM diagnosis is less than 18 months and fewer than 7% of patients exhibit longterm survival greater than 5 years [2; 3; 4; 5]. Currently, the treatment options for GBM include surgical resection, radiotherapy, and temozolomide-based chemotherapy [6; 7]. Despite these treatment availability, the median survival has only been improved from less than 10 months in the late 1970s [8; 9] to approximately 15 months [2; 3; 7; 10]. However, it remains that a more thorough understanding of GBM-centric biology is crucial in the improving clinical outcomes for GBM patients. Therefore, it is critical for early GBM detection and prevention for reducing the high mortality rate. A better understanding of the molecular basis of GBM formation and the identification of markers are essential for the development of preventive therapies targeting the specific GBM-promoting factors and thereby improve prognosis.

Recent studies have identified the role of altered cellular metabolism as a key hallmark of GBM playing an important role in enhancing the invasiveness of tumor cells. Metabolic reprogramming allows invasive cells to generate the energy necessary for colonizing surrounding brain tissue and adapt to new microenvironments with unique nutrient and oxygen availability. Akin to many other cancers, GBM cells exhibit the Warburg effect, where even in the presence of high levels of oxygen, cancer cells exhibit a strong preference for glycolysis [11; 12]. In GBM, glycolysis has been suggested to correlate with tumor proliferation, invasion, angiogenesis, and chemotherapy/radiotherapy resistance [13; 14; 15]. In addition, glycolysis could shape the tumor microenvironment (TME) and regulate immune and inflammatory responses[16]. Also, literature suggests a shift of cancer therapies away from radiotherapy and towards genetic or chemical interventions. Specifically, inhibitors with targets related to glycolysis including hexokinase 2 (HK2) and lactate dehydrogenase A (LDHA) are useful in selectively killing cancer cells [17]. Further, recent work suggests that it is possible to force the differentiation of glioblastoma to astrocytes by shifting the energy preference of the GBM cells back to oxidative phosphorylation [18]. While glycolysis acts as the dominant source of growth substrate in GBM, the role of other metabolic from pathways including tricarboxylic acid (TCA) cycle, oxidative phosphorylation (OXPHOS) for ATP production and pentose phosphate pathway (PPP) which generates the cellular reductant NADPH and macromolecules (nucleotides, amino acids, and fatty acids) have not been completely elucidated. There is limited data available on altered gene expression of the enzymes involved in these metabolic pathways *in vivo*. Thus, a deeper understanding of metabolic changes including glycolysis could be an important step towards the individualized treatment of GBM.

In this study, we investigated the expression level of enzymes required for the glycolytic and mitochondrial metabolism in brain samples from patients with LGG and GBM available from two open-source data sets. We profiled the hallmark gene sets in 174 samples from GBM patients with whole mRNA expression data from REpository for Molecular BRAin Neoplasia DaTa (REMBRANDT) database and compared with 28 normal brain tissues. We also compared genes associated with glycolysis, TCA cycle, PPP and OXPHOS in 530 samples from LGG with 152 samples of GBM patients from the Cancer Genome Atlas (TGGA). Notably, the glycolysis-related risk signature could independently identify patients in the high risk group with poor prognosis.

## 2. MATERIALS AND METHODS

### Data Sets Review

Analysis of differential gene expression in GBM tumors compared with healthy tissues and survival comparisons between cohorts of glioma patients were performed using publicly available datasets available through the REpository for Molecular BRAin Neoplasia DaTa (REMBRANDT) [19; 20; 21]. We selected this dataset (GSE108476) due to its large size and comprehensive information on tumor grading. This dataset was accessed through Georgetown Database of Cancer (G-DOC ®) [22] (study id: 580). This dataset includes information on 671 patients submitted from 14 institutions between 2004 and 2006 [19]. Total RNA from each sample was processed with the Affymetrix HG U133 V2.0 Plus gene expression chips [20]. From this dataset, we extracted data on GBM (N^GBM^ = 174) patients who had both gene expression and survival time available, and astrocytoma (N^ASTRO^ = 116) patients who had both gene expression and survival time available, and healthy (N^NORM^ = 28) patients which had gene expression data available. Gene expression was processed with MAS5 normalization on G-DOC® and extracted for further analysis.

Analysis of differential gene expression in GBM tumors compared with lower grade glioma (LGG) tissues were performed using publicly available datasets available through The Cancer Genome Atlas (TCGA) [23]. We selected these datasets due to their extensive normalization procedure, RNA Seq V2 RSEM, which allows for the comparison of relative gene expression across sample runs [24; 25]. The TCGA-GBM dataset (GSE83130) includes information on 543 patients with World Health Organization (WHO) grade IV gliomas. We extracted GBM patients with total RNA expression available (N^TCGA-GBM^ = 152). The TCGA-LGG dataset includes information on 530 patients with WHO grades II or III gliomas including 134 Oligoastrocytoma, 130 Anaplastic Astrocytoma, 120 Oligodendroglioma, 78 Anaplastic Oligoastrocytoma, 67 Astrocytoma, and 1 Diffuse Glioma patient. We extracted LGG patients with total RNA expression available (N^TCGA-LGG^ = 530). Both TCGA datasets were downloaded from the cBioPortal for Cancer Genomics (cbioportal.org) [26; 27]. Relative RNA Seq V2 RSEM normalized gene expression data was extracted for further analysis.

### Data Sets Analysis

Differential gene expression between normal and GBM patients was analyzed by downloading raw data from the REMBRANDT study from National Center for Biotechnology Information (NCBI) using the GSE accession number (GSE108476). Clinical data and reporter IDs for specific genes were accessed using Georgetown Database of Cancer (G-DOC ®) (study id: 580). Each sample was then categorized by tumor grade, survival time, and diagnosis. GBM, astrocytoma, and non-tumor samples were extracted for further analysis.

Gene expression was compared between GBM patients and healthy patients. GraphPad was used to perform a non-parametric Mann-Whitney U-test with exact p-values. Significance threshold was assigned at p-value < 0.05.

Relative survival within glioma patients was analyzed based on a combined cohort of GBM (N^GBM^ = 174) and astrocytoma (N^ASTRO^ = 116) patients for a combined cohort of glioma (N^GLIOMA^ = 290) patients. Within each gene, the glioma cohort was separated into two groups based on the median value of gene expression. Patients having higher gene expression than the median were assigned to the ‘High Expression’ group. Patients having lower gene expression than the median were assigned to the ‘Low Expression’ group. GraphPad was used to generate Kaplan Meier plots and calculate significance using the Log-rank (Mantel-Cox) test. Significance was assigned at p-value < 0.05.

## 3. RESULTS

A salient feature of GBM cells is that they adjust their metabolic profile to fulfill the bioenergetics and anabolic demands of the high rates of proliferation [14; 28]. Yet, little is known about the metabolic changes at stages of the disease. To have a complete overview of the metabolic genes expressed in LGG and GBM stages of disease, we analyzed the transcription profiling of enzymes involved in glycolysis, PPP, TCA, and OXPHOS in brain samples from patients with LGG and GBM.

### Key Glycolysis-related and PPP gene expression Increases in GBM Patients

The glycolytic pathway is a well-orchestrated and characterized series of ten enzymatic reactions converting glucose to pyruvate (***Figure 1A***). We evaluated the mRNA expression levels of these ten enzymes. These ten genes include HK2 (hexokinase 2), GPI (glucose-6-phosphate isomerase), PFK* (6-phosphofructokinase), ALDOC (aldolase, fructose bisphosphate C), GAPDH (glyceraldehyde 3-phosphate dehydrogenase), PGK1 (phosphoglycerate kinase 1), PGAM1 (phosphoglycerate mutase 1), ENO1 (enolase 1), PKM1/2 (pyruvate kinase M1/2), and LDHA (lactate dehydrogenase A). As shown in ***Figure 1B*,** expression of the rate limiting glycolytic transcript, HK2 (p < 0.0001), was significantly increased in GBM patients compared to normal patients. The HK2 overexpression has been shown to contribute to the enhanced glycolytic rate and tumor progression and confer resistance to apoptosis to cancer cells [29]. Along with this, a majority of glycolytic transcripts including ALDOC (p < 0.0001), GAPDH (p < 0.001), PGK1 (p < 0.05), and PGAM1 (p < 0.0001), were significantly increased in GBM patients compared to normal patients. ALDOC is specific to the brain region and is involved in the fructolysis process which is shown to promote the Warburg effect by preferentially downregulating OXPHOS and mitochondrial respiration and increase aerobic glycolysis that may aid metastases that initially have low oxygen supply [13; 30]. GAPDH has been shown to enhance mitophagy and induce cancer cell survival [31]. High expression of PGK1 is correlated with tumor proliferation, metastasis, occurrence, development and prognosis prediction [32; 33]. Overexpression of PGAM1 is linked with tumor growth, survival, and invasion in several cancers including GBM [34; 35]. Expression of LDHA and GPI was significantly reduced in GBM samples compared to normal patients (p < 0.0001). This is interesting as LDHA enzyme converts pyruvate into lactate generating NAD+ and diverting glycolysis-derived pyruvate from the mitochondrial oxidative pathway [36]. As a result, this decreased expression and activity of LDHA would favor the routing of pyruvate into mitochondria where it can be further metabolized through TCA and oxidative phosphorylation. GPI catalyzes G6P to F6P and decrease in GPI will interrupt the glycolytic flux by increasing the intracellular G6P pool, however it also could reactivate PPP for the cells’ energy needs. In contrast, mRNA expression of PFKL (p = 0.62), ENO1 (p = 0.46), and PKM1/2 (p = 0.17) showed no significant changes.

**Figure 1.**
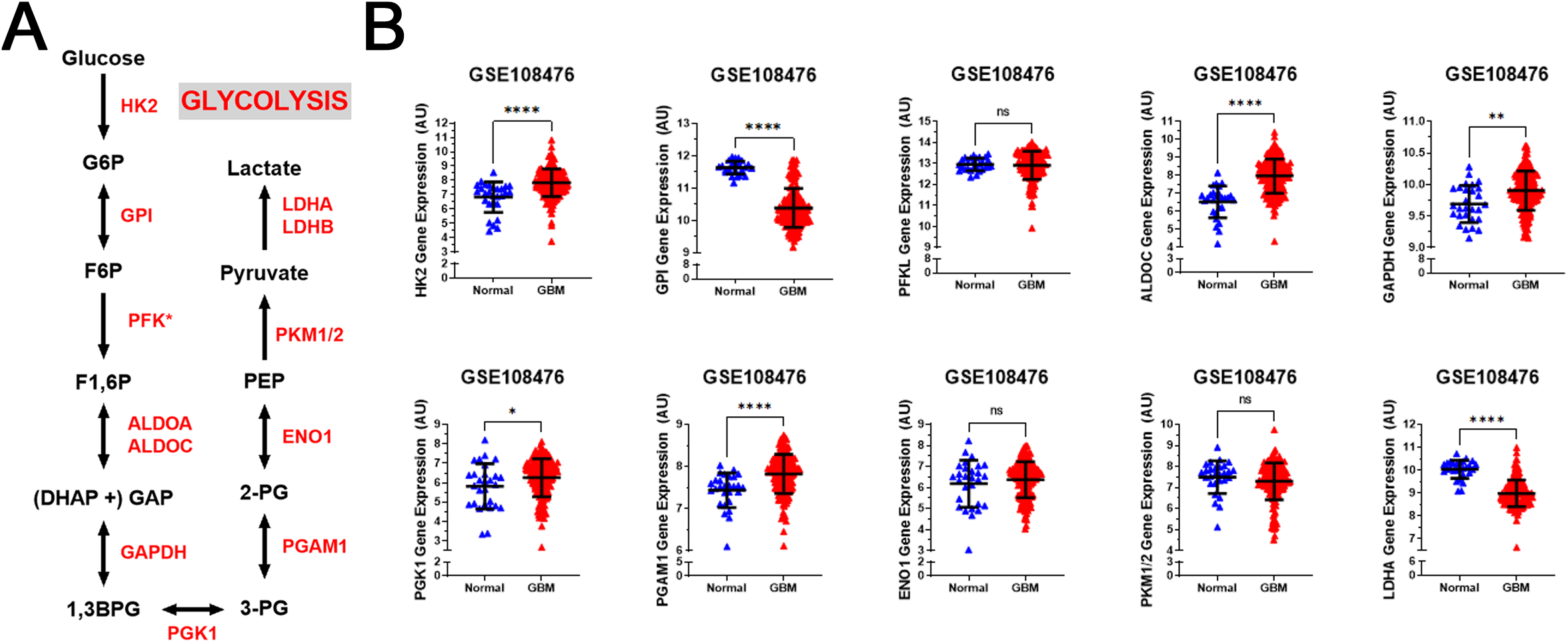
Glycolytic genes are significantly overexpressed in GBM compared with normal tissue. (**A**) Schematic overview of glycolysis. Abbreviations of the enzymes are as follows: hexokinase 2 (HK2), glucose-6-phosphate isomerase (GPI), phosphofructokinase liver isoform (PFKL), aldolase A (ALDOA), glyceraldehyde 3 phosphate dehydrogenase (GAPDH), phosphoglycerate kinase 1 (PGK1), phosphoglycerate mutase 1 (PGAM), enolase 1 (ENO1), and pyruvate kinase M2 (PKM2), lactate dehydrogenase (LDH). Abbreviations of the metabolites are as follow: glucose-6-phosphate (G6P), fructose-6-phosphate (F6P), fructose 1,6-biphosphate (F1,6BP), glyceraldehyde 3-phosphate (GAP), and dihydroxyacetone phosphate (DHAP), 1,3-biphosphoglycerate (1,3BPG), glycerol-3-phosphate (3-PG), glycerol-2-phosphate (2-PG), phosphoenolpyruvate (PEP) (**B**) Relative gene expression of different glycolytic enzymes in the clinical data set GSE108476 consisting of GBM brain tissues (n= 174) and normal (n= 28) brain tissues. Gene expression data from the Repository for Molecular Brain Neplasia Data (REMBRANDT) via Affymetrix HG U133 v2.0 Plus. Significance calculated via Mann-Whitney Test. * indicates p < 0.05, ** indicates p < 0.001, **** indicates p < 0.0001, N/S indicates not significant.

We also examined the relative gene expression of enzymes involved in the oxidative phase of PPP. PPP branches from glucose 6-phosphate (G6P), produces NADPH and ribose 5-phosphate (R5P), and shunts carbons back to the glycolytic or gluconeogenic pathway (***Figure 2A***). We found that two of PPP genes, 6-phosphogluconolactonase (PGLS) and 6-phosphogluconate dehydrogenase (PGD) were significantly upregulated (p < 0.0001) in GBM patients compared to healthy patients while expression of glucose 6-phosphate dehydrogenase (G6PD) was found to be significantly downregulated (p < 0.001) in GBM patients than in healthy patients (***Figure 2B***). The decrease in G6PD which is the rate limiting enzyme for PPP indicates that the possibility of glucose metabolism shift from PPP to neuronal glycolysis leading to higher susceptibility to oxidative stress in the brain. The increase in PGLS and PGD indicates that the production of R5P required for nucleotide synthesis during cell growth and NADPH production for energy requirements. Together, these results imply an overall increase in glycolytic and PPP genes driving the production of excess ATP and nucleotides which give way to unchecked proliferation in GBM cells.

**Figure 2.**
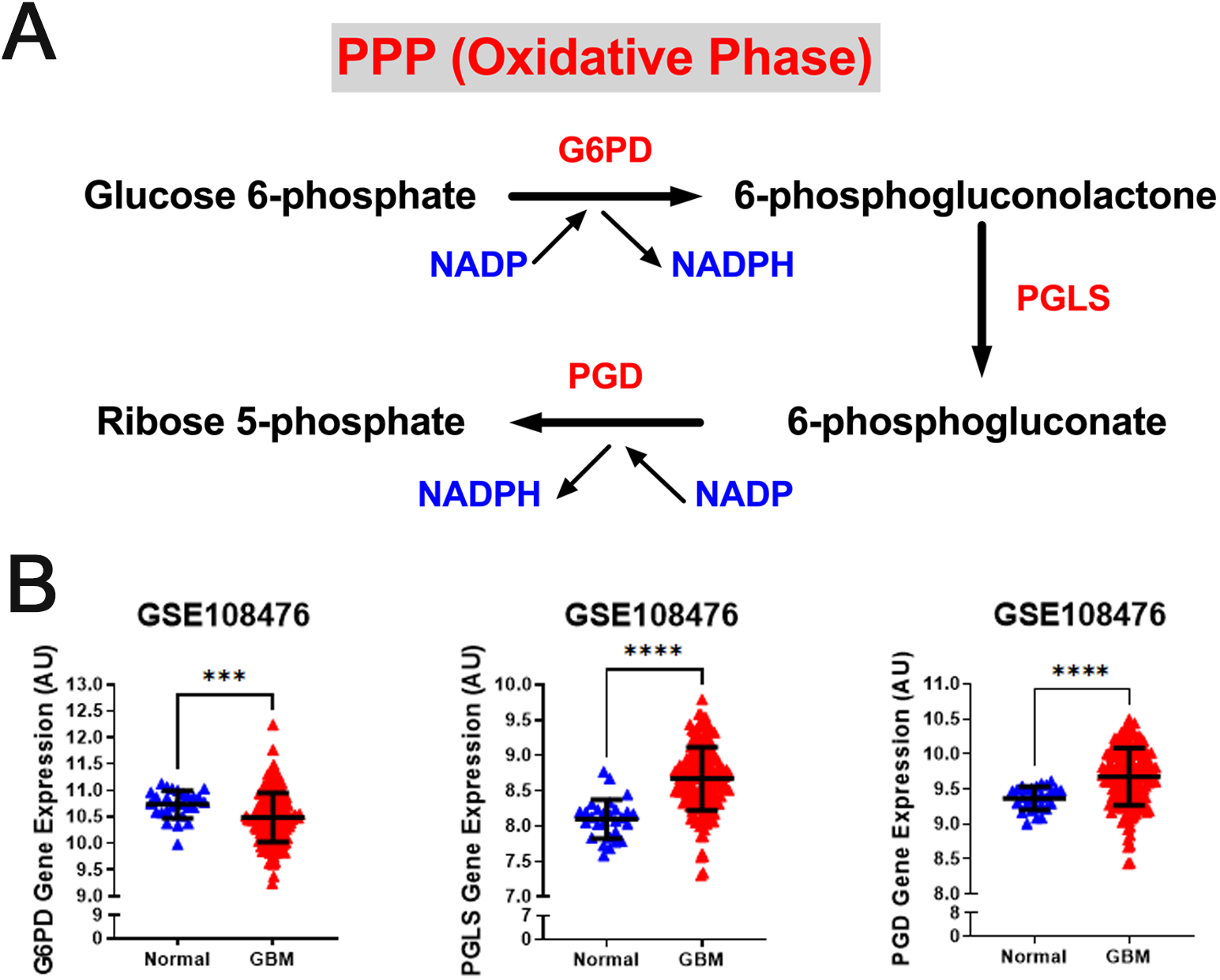
Gene expression of PPP enzymes are significantly overexpressed in GBM compared with normal tissue. (A) Schematic overview of the oxidative phase of the pentose phosphate pathway. Abbreviations of the enzymes are as follows: glucose-6-phosphate dehydrogenase (G6PD), 6-phosphogluconolactonase (PGLS), 6-phosphogluconate dehydrogenase (PGD). Activation of the two dehydrogenase enzymes, G6PD – the rate-limiting enzyme – and PGD, results in the production of NADPH, H+ ions, and ribose 5-phosphate. (B) Relative gene expression of G6PD, PGLS, and PGD in the clinical data set GSE108476 consisting of GBM brain tissues (n= 174) and normal (n= 28) brain tissues. Each are upregulated in cancer tissue compared with normal brain tissue. Gene expression data from the Repository for Molecular Brain Neplasia Data (REMBRANDT) via Affymetrix HG U133 v2.0 Plus. Significance calculated via Mann-Whitney Test. *** indicates p < 0.001, **** indicates p < 0.0001.

### Gene expression of Enzymes of TCA Cycle Decreases in GBM Patients

While normal cells utilize glucose as the main source of pyruvate entering the TCA cycle, tumor cells often shunt glucose away from the TCA cycle for catabolism through anaerobic glycolysis. Also, inherited and acquired alteration of TCA cycle enzymes have been demonstrated in different cancers [37; 38]. We examined the gene expression levels of three key metabolic enzyme of TCA cycle: pyruvate dehydrogenase E1 alpha 1 subunit (PDHA1), isocitrate dehydrogenase (NADP(+)) 2, mitochondrial (IDH2), and succinate dehydrogenase (SDHB). We observed that all three genes were significantly lower than that in normal patients (p < 0.0001) (***Figure 3A***). Loss of PDHA1 has been shown to decrease mitochondrial OXPHOS and promote aerobic glycolysis in tumor cells and promotes Warburg effect [39; 40]. SDH catalyzes the oxidation of succinate to fumarate in the TCA cycle while simultaneously reducing ubiquinone to ubiquinol in the mitochondrial electron transport chain (ETC) reactions [41]. Lack of SDHB has been shown to impair mitochondrial oxygen consumption and commit cells to ferment glucose for sustaining their energetic needs [42]. The lack of SDH In GBM cells possibly results in a complete deficiency of the enzyme activity and abnormal accumulation of succinate in GBM. Isocitrate dehydrogenases 2 (IDH2) catalyzes isocitrate to alpha-ketoglutarate and generates NADPH in the TCA cycle. Loss of IDH2 has been shown to impair oxidative bioenergetics, elevate reactive oxygen species (ROS) production, and promote exaggerated mitochondrial dynamics in prostate cancer cells [43]. Our observations are consistent with numerous studies showing that decreased levels of the TCA associated genes in GBM promotes the Warburg effect and tumor growth.

**Figure 3.**
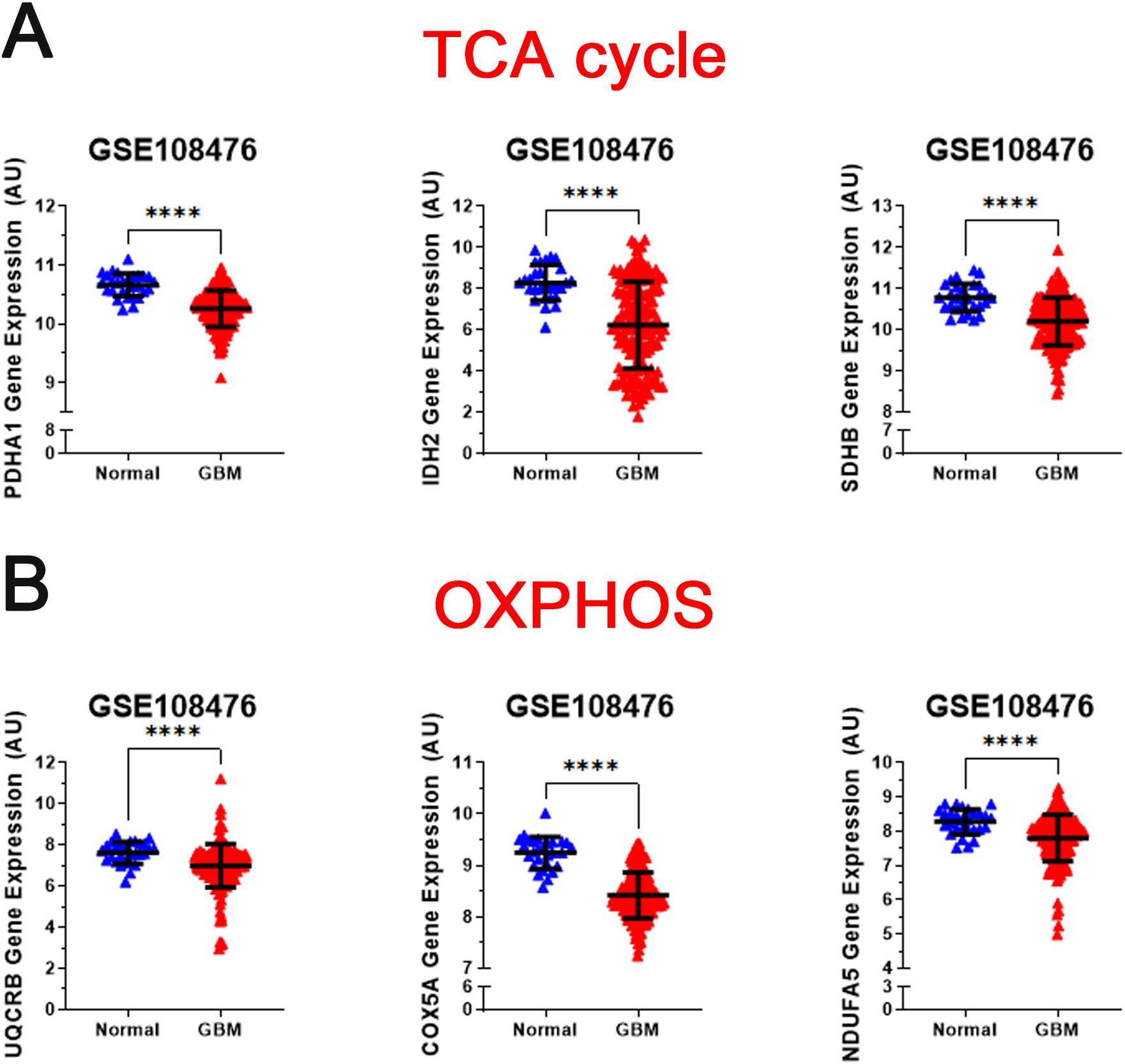
Analysis of gene expression of oxidative mitochondrial metabolism. (A) Relative gene expression of PDH1, IDH2, and SDHB implicated in the TCA cycle in the clinical data set GSE108476 consisting of GBM brain tissues (n= 174) and normal (n= 28) brain tissues. PDH1, IDH2, and SDHB gene expression is significantly under expressed in GBM tissue compared with normal tissue. (B) Relative gene expression of UQCRB, COX5A, and NDUFA5 implicated in oxidative phosphorylation in the clinical data set GSE108476 consisting of GBM brain tissues (n= 174) and normal (n= 28) brain tissues. UQCRB, COX5A, and NDUFA5 are significantly under expressed in GBM tissue when compared to normal tissue. Gene expression data from the Repository for Molecular Brain Neplasia Data (REMBRANDT) via Affymetrix HG U133 v2.0 Plus. Significance calculated via Mann-Whitney Test. *** indicates p < 0.001, **** indicates p < 0.0001.

### Gene expression of Enzymes of Mitochondrial Oxidative Metabolism Decreases in GBM Patients

The regulation of the TCA cycle is cued in with its constant feedback with OXPHOS is critical for metabolic activities in normal cells. In contrast, in tumor cells glycolysis is enhanced and OXPHOS capacity is reduced in tumor cells [44; 45]. To test whther this was the case for GBM, we analyzed the expression profile of representative genes involved in the formation of the electron transport chain complexes: NADH: Ubiquinone Oxidoreductase Subunit A5 (NDUFA5 -complex I); Ubiquinol-Cytochrome C Reductase Binding Protein (UQCRB-complex III), Cytochrome C Oxidase Subunit 5A (COXA5 (COX5A-complex IV). As shown in ***Figure 3B*,** all three genes were significantly reduced (p < 0.0001) in GBM patients compared with healthy patients. This corroborates with the changes observed in glycolysis and TCA cycle genes.

Altogether, these results indicate that in GBM patients the expression of glycolytic related genes is higher relative to normal brain, while the expression of TCA cycle and OXPHOS genes are reduced compared to normal brain (***Figures 1–3***). Importantly, Niclou and colleagues have investigated the importance of glycolytic enzymes in GBM adaption and survival including HK2, ALDO, and PGAM1 in 5 patient-derived GBM stem-like cell lines and found that inhibition of these glycolytic genes pathway strongly affects GBM growth in patient-derived intracranial mouse models [46]. This comprehensive study on glycolytic enzymes are in agreement line with our analyses whereby changes in transcript expression of glycolysis, TCA cycle and OXPHOS, in GBM are consistent with a Warburg-type metabolism.

### Expression of Glycolytic Genes Correlates with Poor Patient Survival in GBM

We next investigated the survival correlation of GBM patients with key regulators of aerobic glycolysis and mitochondrial metabolism using the REMBRANDT dataset (GSE108476). This data was analyzed via Kaplan-Meier plots and significance was determined with a Log-rank (Mantel-Cox) test (***Figure 4***). In two rate limiting steps of glycolysis, the overall survival rate was significantly lower in high expression of HK2 (p = 0.0017) and PKM2 (p = 0.0079) (***Figure 4A***). There was not a significant correlation found within the two remaining rate limiting enzymes, PFKL and LDHA. Interestingly, the opposite trend was noted within the expression of genes associated with the PPP. Low expression of G6PD (p = 0.0196) and PGLS (p = 0.0494) was correlated with worse patient outcomes (***Figure 4B***). There was no significant difference in the survival of patients with PGB gene expression. In the TCA cycle-related genes, lower expression of SDHB (p < 0.0001) correlated to worse patient outcomes while PDHA1 and PGD displayed no significant difference in survival (***Figure 4C***). Finally, in the OXPHOS genes, we found a significant correlation with lower expression of COX5A (p < 0.0001) and poor clinical outcomes, while UQCRB and NDUFA5 showed no significant correlation (***Figure 4D***). Taken together, this indicates that patients’ gene expression profiles could be correlated with overall clinical outcome. In particular, patients with higher expression of glycolytically related genes have poor prognosis and survival rates indicating a potential route for therapeutic development.

**Figure 4.**
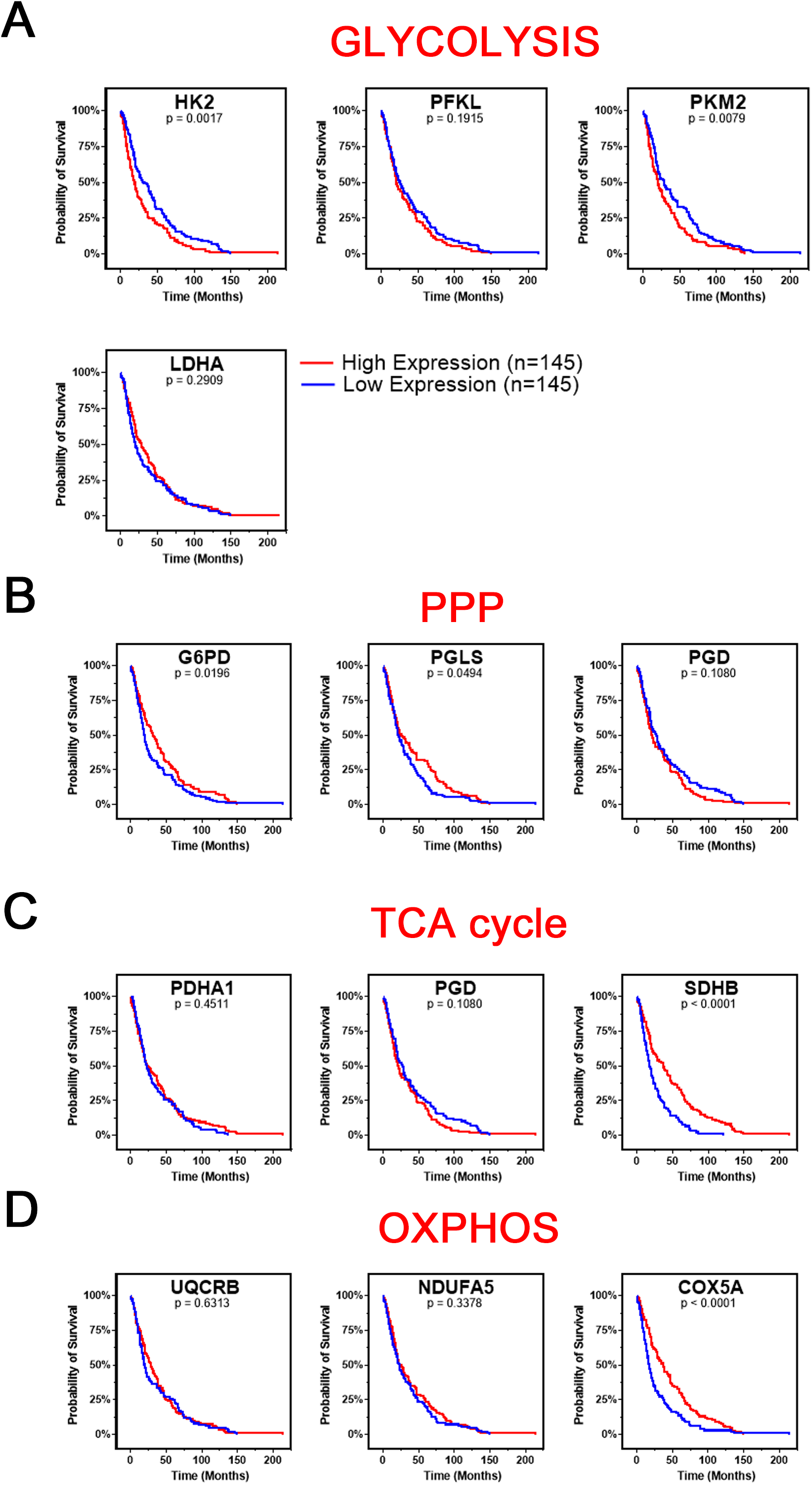
Changes in gene metabolic gene expression is associated with poor patient prognosis. Shown are the Kaplan-Meier overall survival curves of GBM patients according to the designated gene expression levels above or below the median value based. (A) High expression of glycolysis related genes, HK2 and PKM2, correlated with poor overall GBM patient’s survival. There was no significant difference in survival based on PFKL or LDHA expression. (B) Low expression of PPP related genes, G6PD and PGLS, correlated with poor overall GBM patient’s survival. There was no significant difference in survival based on PGD expression. (C) Low expression of TCA related gene, SDHB, correlated with poor overall GBM patient’s survival. There was no significant difference in survival based on PDHA1 or PGD expression. (D) Low expression of OXPHOS related gene, COX5A, correlated with poor overall GBM patient’s survival. There was no significant difference in survival based on UQCRB or NDUFA5 expression. Gene expression in GBM tumors and associated patient survival from the REpository for Molecular BRAin Neoplasia DaTa (REMBRANDT) (GSE108476) via Affymetrix HG U133 V2.0 Plus gene expression chips. Gene expression was processed with MAS5 normalization on G-DOC® before comparison. Significance calculated via Log-rank (Mantel-Cox) test. Low expression, n = 145. High expression, n = 145. Exact p-values reported.

### Gene Expression of Enzymes Involved in Glycolysis, TCA Cycle, Mitochondrial Oxidative Metabolism, and PPP is Higher in GBM compared with LGG

Brain lower-grade and intermediate-grade glioma (LGG-classified as grade I, II, or III) are infiltrative neoplasms that has highly variable clinical behavior but often poor prognosis [47]. While phenotypically less aggressive than GBM, LGGs represent a disease in need of further exploration to advance treatment options and prolong patient survival. To better understand the metabolic alterations in LGGs in comparison to GBMs, we examined the expression of glycolytic transcripts and transcripts of PPP, TCA cycle and OXPHOS in a clinical data set with 152 GBM and 530 LGG patients from the TCGA database (***Figure 5***). Compared with LGG, GBM tumors expressed higher levels of LDHA (8.1 fold) and genes involved in glucose uptake including HK2, PFKL, and PKM (3.7, 2.1, 2.9 fold) (***Figure 5A***). Interestingly, all ten glycolytic related genes were significantly higher expression in GBM compared to LGG samples (p < 0.0001). Also, GBM tumors exhibited elevated expression of PPP genes including G6PD, PGLS, and PGD (p < 0.0001) (***Figure 5B***). In addition, GBM tumors expressed higher levels of TCA cycle genes including IDH2 (the primary producer of NADPH in GBM) [48] compared to LGG (***Figure 5C***). The GBM-activated PPP also generate NADPH, which is a reducing equivalent for tumor cells affected by the Warburg effect [49]. Interestingly, LDHA was the highest fold change in GBM compared to LGG indicating that the conversion of pyruvate to lactate and Warburg related properties were exacerbated during the tumor shift. Together, these results suggest that GBM tumors metabolically rely on the Warburg effect.

**Figure 5.**
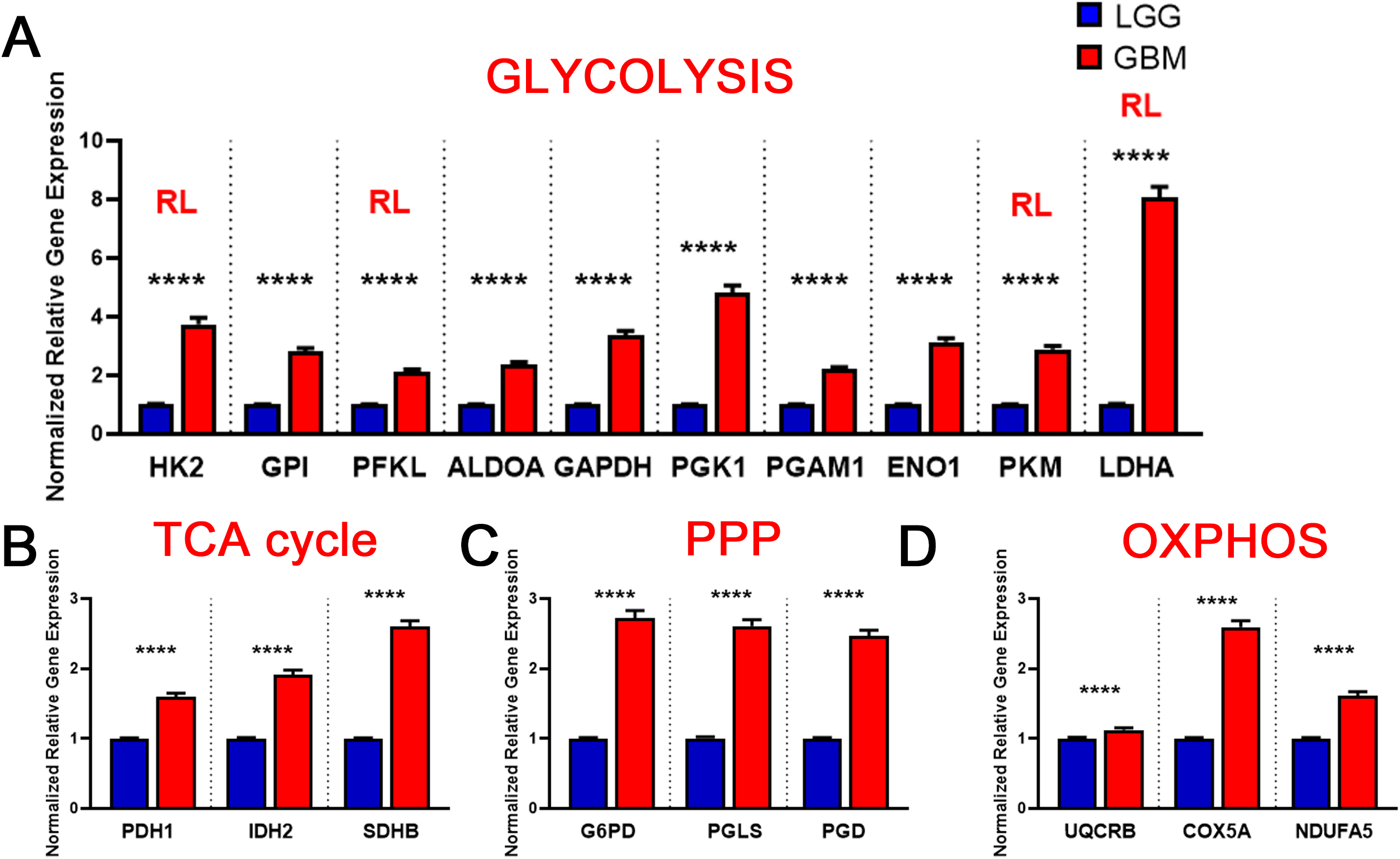
Metabolic gene expression higher in GBM patients than in LGG patients. (A) Higher glycolytic gene expression in glioblastoma (GBM) patients compared with lower grade glioma (LGG) patients. Red **RL** indicates association with a rate limiting step of glycolysis. (B) Higher relative gene expression of TCA cycle genes in glioblastoma (GBM) patients compared with lower grade glioma (LGG) patients.(C) Higher relative gene expression of PPP genes in glioblastoma (GBM) patients compared with lower grade glioma (LGG) patients.(D) Higher relative gene expression of oxidative phosphorylation (OXPHOS) genes expression in glioblastoma (GBM) patients compared with lower grade glioma (LGG) patients. Gene expression data from The Cancer Genome Atlas (TCGA) including TCGA-LGG and TCGA-GBM (GSE83130) downloaded from the cBioPortal for Cancer Genomics. Relative RNA Seq V2 RSEM normalized gene expression was used for analysis. Significance calculated via Mann-Whitney U Test. LGG n=530, GBM n=152 **** indicates p<0.0001.

**Figure 6.**
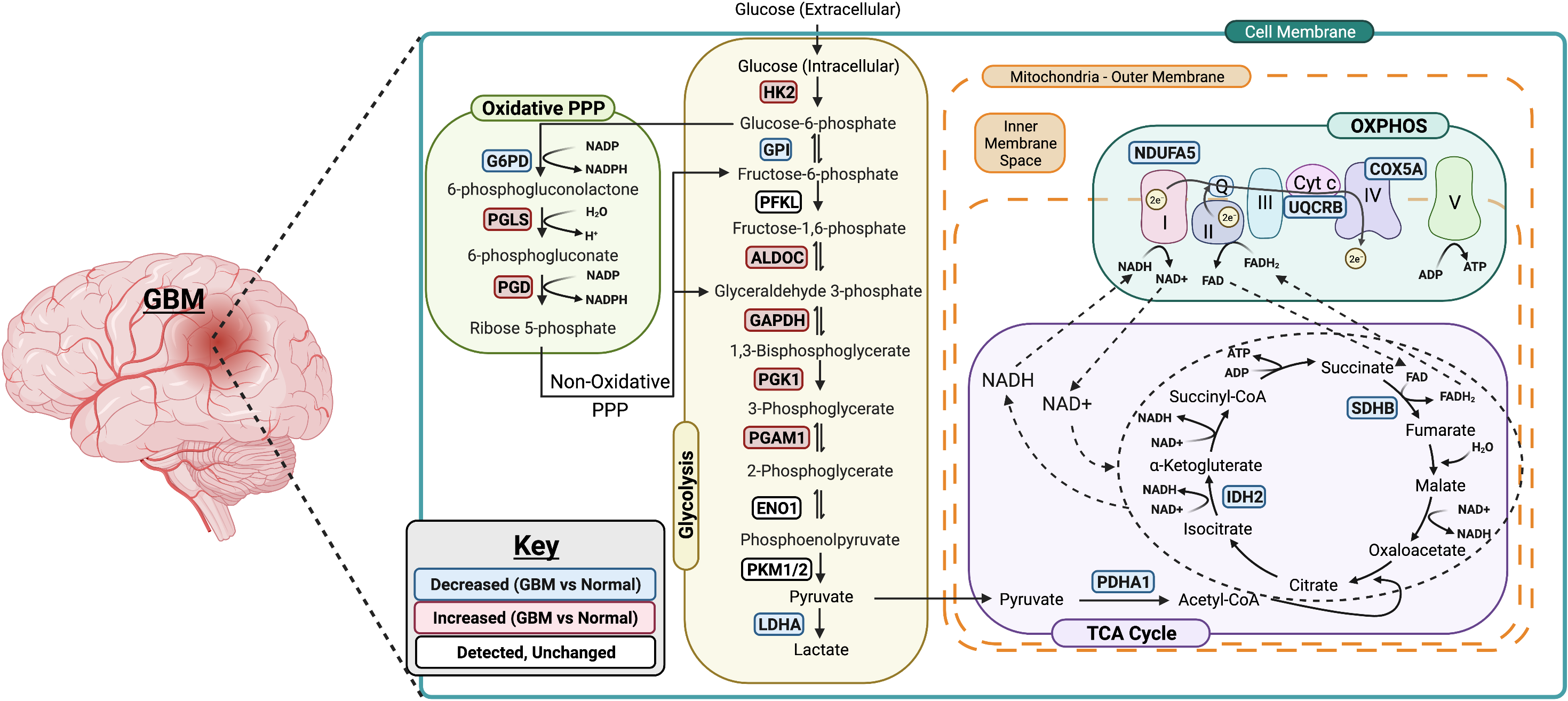
Summary of metabolic changes in GBM. A simplified model showing gene changes of metabolic enzymes during the progression of GBM. One of the most pronounced transcriptional changes during the progression of GBM is the increase in genes associated with rate limiting enzymes in glycolysis including HK2, ALDOC, GAPDH, PGK1 and PGAM1. This allows glucose to enter in the cells and be converted to pyruvate. The PPP-associated genes are upregulated as well suggesting the entrance into oxidative phase of PPP. Surprisingly, the genes associated with TCA cycle and OXPHOS are downregulated in GBM which suggests that GBM primarily relies on glycolysis for its energy needs. Figure was created using Biorender.

Similarly, we evaluated the transcripts levels of representative OXPHOS genes-UQCRB, COX5A, and NDUFA5-in LGG and GBM patients. The expression of OXPHOS-related genes were significantly higher in GBM compared to LGG (p < 0.0001) (***Figure 5D***). This result suggest that, in contrast to a general increase in glycolysis genes, TCA and OXPHOS genes remained higher in GBM. Compared to normal patients, GBM patients expressed lower levels of genes involved in TCA cycle and OXPHOS, but at significantly higher levels than those found in LGG (***Figure 5 C,D***).This indicates that GBM tumors generate ATP via OXPHOS as well compared to LGG under aerobic conditions. Together, these results suggest that LGG tumors metabolically rely primarily on the Warburg effect while GBM utilizes TCA cycle and OXPHOS as well for their high energy demand.

## 4. DISCUSSION

This study represents a genomic analysis to delineate the metabolic foundations of GBM and conclude that the genetic changes in the glycolytic genes was more reflective of the disease aggressiveness and prognosis. Specifically, we report that the changes in transcript expression of glycolysis and PPP genes in GBM is consistent with a Warburg-type metabolism and show a positive correlation between glycolytic gene expression and the risk of developing GBM and poor patient survival. We found that the metabolic trends in GBM varies significantly from LGG especially with OXPHOS and TCA cycle related genes. GBM tumors displayed distinct Warburg-like genetic features, with increased HK2, PGAM1, ALDOC, and PGK1 expression that facilitates lactate production and confers resistance to hypoxic stress in cancer cells [32; 50]. The Warburg effect drives the biosynthesis of nucleotides, lipids, and proteins to support rapid cell proliferation, as well as the disruption of tissue architecture to facilitate tumor motility [49]. NADPH is a key component in this process, thus the elevated PPP, in GBM tumors provide additional support implying a pathogenic role of glucose metabolism for this disease.

Although, it is now well accepted that the aerobic glycolytic phenotype is associated with an impaired mitochondrial oxidative metabolism [51], recent studies have indicated a new paradigm that both glycolytic and mitochondrial metabolism are used by cancer cells for ATP production and macromolecule synthesis [52; 53; 54]. In concordance with these studies, our analysis in GBM patients show high expression levels of genes related to mitochondrial metabolism compared to LGG patients. The fact that in GBM the expression levels of TCA and OXPHOS genes are higher than those in LGG patients suggests the metabolic shift in GBM cells when progressing from LGG to GBM. GBM have high expression of HK2, ALDOC, GAPDH, PGK1, and PGAM1 as well as display a significant increase in PGLS and PGD expression while low expression of TCA cycle and OXPHOS related genes compared to healthy patients. It appears that metabolic readjustments (that is, glycolysis shift) occurs when the transition from LGG to GBM occurs as genes related to glycolysis, PPP, TCA cycle and OXPHOS are higher in GBM compared to LGG. Overall, these results begs the need to evaluate the differential signature of these metabolic regulators in normal, LGG and GBM patients and delineate the metabolic transition especially from the less aggressive LGG to GBM state. Further studies such as protein expression analysis are therefore required to understand the specific mechanisms underlying the metabolic changes observed in LGG and GBM, although it might be difficult due to shortage of human GBM samples available.

Our analysis of clinical outcomes showed that GBM patients with high expression of HK2 and PKM2 in the glycolysis related genes had shorter overall survival providing the evidence that HK2 and PKM2 constitutes important targets for GBM therapy. Knockdown of HK2 gene has been shown to strongly inhibit GBM growth indicating that HK2 is essential for GBM growth and also impact overall survival [55; 56]. The switch in splice isoforms from the adult pyruvate kinase muscle 1 (PKM1) to the fetal PKM2 has been attributed to promote aerobic glycolysis and tumor growth in lung cancer cell lines [57; 58]. Interestingly, the low expression of two metabolic genes involved in mitochondrial metabolism-SDHB and COX5A related to TCA cycle and OXPHOS, respectively, was associated with poor patient overall survival which parallels their significant decreases in GBM samples compared to normal brain. Succinate dehydrogenase (SDHB, also known as mitochondrial respiratory chain complex II) is a key respiratory enzyme located on the inner mitochondrial membrane, which links the TCA cycle with OXPHOS and plays the important roles in both TCA cycle and OXPHOS [59]. The low expression of SDHB has been shown to promote aerobic glycolysis [60], and the lack of SDHB function has been implicated in the occurrence and development of multiple kinds of tumors, including liver cancer, renal cancer and colorectal cancer [60; 61]. COX5A, one of the three mitochondrial-encoded subunit of COX, is the terminal enzyme of the respiratory chain and key regulator of OXPHOS [62]. It has been shown that HK2 inhibits OXPHOS and promotes tumor proliferation [56]. Our survival data implies that the overall survival is poor in GBM patients where aerobic glycolysis occurs along with active suppression of OXPHOS. Prevention of GBM is an unmet medical need and current clinical interventions do not extent the survival rate beyond 15 months. Interestingly, our data suggests that high expression levels of HK2, and PKM2 appear to have important clinical implication for patients with GBM and in agreement with studies demonstrating that inhibition of glycolysis improves survival in GBM [63]. By contrast, low gene expression of SDHB (TCA cycle) and COX5A (OXPHOS) was associated with an increased risk of GBM. This suggests that expression of these glycolytic and metabolic enzymes could be used as a new biomarker for the risk of developing GBM.

Altered metabolic pathways are one of the hallmarks of cancerous cells including GBM. While plethora of studies have attributed numerous pathways that enhance the tumor cells invasiveness in the brain microenvironment as the primary driving forces behind GBM, recent studies have identified a role for cellular metabolism reprogramming in GBM invasion. Metabolic reprogramming acts as the energy source necessary for invasive cells to colonize surrounding brain tissue and adapt to new microenvironments with unique nutrient and oxygen availability. Historically, enhanced glycolysis, even in the presence of oxygen (the Warburg effect) has dominated GBM research with respect to tumor metabolism. While this study allowed us to dive slightly deeper into metabolic pathways beyond glycolysis associated with these metabolic changes, it is important to note that glycolysis and mitochondrial metabolism likely mark merely two of the dozens of pathways affected during the process of GBM progression. There are also several limitations to our study. First, the clinical tumor samples analyzed in our study were limited to those of REMBRANDT and TCGA. This is due to the small amount of public LGG datasets that span multiple the multiple modalities necessary for our study (eg, RNA-seq, clinical/survival). To this end, it may be interesting to explore whether glycolytic and mitochondrial oxidative metabolism gene expression correlates with patient overall survival based on different phases and mutations of LGG and GBM in a larger multimodal dataset. Second, while our results revealed consistently robust statistical associations, they do not imply a direct cause and effect relationship between LGG progression to GBM, glycolysis, and metabolism expression. Nevertheless, as shown in the work of Niclou and colleagues, the direct effect of key glycolytic enzymes revealed results corroborating with our *in silico* analysis [46]. Furthermore, although the purpose of our study was to identify clinically associated genomic changes in glycolysis and metabolic genes in human GBM, our results extend previous *in vitro* and *in vivo* findings to show relevance in human tumor data. This may therefore encourage mechanistic exploration of metabolic genes in glycolysis, PPP, TCA cycle and OXPHOS induced changes in tumor aggressiveness and invasiveness contributing to patient survival outcomes. To this end, recent global profiling experiments have identified roles for lipid, amino acid, and nucleotide metabolism in tumor growth and invasion [14; 64]. A thorough understanding of the metabolic traits that define invasive GBM cells may lead to novel therapeutic targets for this devastating disease.

In summary, our analysis has expanded our knowledge of the glioma metabolic alteration landscape, emphasized the relevance of metabolic changes particularly glycolysis-related genes as a modality for clinical outcomes and survival prediction of GBM patients. Combined, these findings are an important step forward in our understanding of the role of metabolic reprogramming in glioma as drivers of the tumor and could be potential prognostic targets in GBM therapies. Our findings may provide novel insights for GBM research and guidance for individual therapy.

## Author Contributions

Conceptualization, S.K; methodology, K.M.S. and S.K.; data analysis, K.M.S., C.W., and S.K.; writing—original draft preparation, K.M.S., C.W., and S.K.; writing—review and editing, K.M.S., C.W., and S.K. All authors have read and agreed to the published version of the manuscript.

## Acknowledgements

The results shown here are in whole or part based upon data generated by the TCGA Research Network: https://www.cancer.gov/tcga

The data utilized in this study was provided in part by the Georgetown Database of Cancer (G-DOC®), a project of the Georgetown Lombardi Comprehensive Cancer Center designed to provide advanced translational research tools to the scientific community.

